# Temperature but not the circadian clock determines nocturnal carbohydrate availability for growth in cereals

**DOI:** 10.1101/363218

**Authors:** Lukas M. Müller, Leonard Gol, Jong-Seong Jeon, Andreas P.M. Weber, Seth J. Davis, Maria von Korff

**Author notes:** Corresponding authors: Lukas M. Müller, Maria von Korff. Other authors’ email addresses: Leonard Gol, Jong-Seong Jeon, Andreas PM Weber, Seth J Davis.

## Abstract

The circadian clock is considered a key target for crop improvement because it controls metabolism and growth in Arabidopsis. Here, we show that the clock gene *EARLY FLOWERING 3* (*ELF3*) controls vegetative growth in Arabidopsis but not in the cereal crop barley. Growth in Arabidopsis is determined by the degradation of leaf starch reserves at night, which is controlled by ELF3. The vegetative growth of barley, however, is determined by the depletion of leaf sucrose stores through an exponential kinetics, presumably catalyzed by the vacuolar sucrose exporter *SUCROSE TRANSPORTER 2* (*SUT2*). This process depends on the sucrose content and the nighttime temperature but not on ELF3. The regulation of starch degradation and sucrose depletion in barley ensures efficient growth at favorable temperature as stores become exhausted at dawn. On cool nights, however, only the starch degradation rate is compensated against low nighttime temperatures, whereas the sucrose depletion rate is reduced. This coincides with reduced biomass in barley but not in Arabidopsis after growth in consecutive cool nights. The sucrose depletion metabolism determines growth in the cereal crops barley, wheat, and rice but is not generally conserved in monocot species and is not a domestication-related trait. Therefore, the control of growth by endogenous (clock) versus external factors (temperature) is species-specific and depends on the predominant carbohydrate store. Our results give new insights into the physiology of growth in cereals and provide a basis for studying the genetics and evolution of different carbohydrate stores and their contribution to plant productivity and adaptation.

**Significance Statement:** The circadian clock controls growth in the model plant *Arabidopsis thaliana* by regulating the starch degradation rate so that reserves last until dawn. This prevents nocturnal starvation until photosynthesis resumes. The cereal crops barley, wheat and rice, however, predominantly consume sucrose instead of starch as carbohydrate source. We find that carbohydrate supply from sucrose at night is regulated by enzyme kinetics and night-time temperature, but not the circadian clock. We postulate that the regulation of growth depends on the predominant carbohydrate store, where starch degradation is controlled by endogenous cues (clock) and sucrose depletion by external factors (temperature). These differences in the regulation of carbohydrate availability at night may have important implications for adapting crops yields to climate change.

## Introduction

Plants, as photoautotrophic organisms, harvest light energy through photosynthesis during the day to drive growth, development, and reproduction. To supply the energy demand during the night, plants partition photosynthetic assimilates for transitory storage in leaves during the day and deplete them at night (1–3). Arabidopsis accumulates almost exclusively starch in its chloroplast during the day and uses the circadian clock to exhaust the starch stores in coincidence with the end of the night (4). This circadian control of starch degradation provides a benefit for growth at night because starch is neither unproductively sequestered nor prematurely depleted until photosynthesis resumes at dawn (1–3). The clock control of starch degradation is maintained in a wide range of conditions, including fluctuating and interrupted photoperiods (4), varying light intensity (5) and low night temperature (6). As a consequence, the circadian control of starch degradation in the chloroplast during the night is a key trait for optimal growth in Arabidopsis (4). However, the cereal crops barley, wheat and rice predominantly accumulate soluble carbohydrates, particularly sucrose, to high amounts in the leaf vacuole during the day and only little amounts of starch in the chloroplast (7–14). Furthermore, barley and wheat, but not rice, can transform abundant sucrose in the vacuole into the storage intermediate fructan during the day to mitigate the osmotic consequences of sucrose accumulation in the leaf cell (7). The conversion between sucrose and fructans is interrelated and determined by the concentration of vacuolar sucrose (7). Due to its molecular size and complex structure, fructans require to be converted back to sucrose before they can be released from the vacuole (7). Therefore, the export of sucrose from the vacuole but not the degradation of crystalline starch in the chloroplast, is the deciding pathway for carbohydrate supply in cereal crops during the night (8, 10, 11, 14, 15). Work from Gordon *et al*. and Farrar *et al*. and others in the 1980s have described carbohydrate fluxes in the barley leaf during the day/night cycle, but it remained unknown if the circadian clock controls depletion of sucrose and starch from the leaf (1, 10, 15–18) and what consequences result for overall growth when sucrose export from the vacuole but not starch degradation in the chloroplast dominates carbohydrate supply during the night (1, 11, 17). The circadian clock is structurally and functionally conserved between Arabidopsis and the cereal crops barley, wheat and rice (19–21), suggesting that the circadian regulation of metabolism and growth as observed in Arabidopsis offers a target for crop improvement (20, 21). However, the link between the circadian clock, carbohydrate metabolism and growth is not well understood in crops. Here, we investigate the link between temporal control of the carbohydrate supply during the night and vegetative growth in the cereals in comparison to Arabidopsis.

## Results and discussion

### The clock component ELF3 determines metabolism and growth in Arabidopsis but not in barley

We measured the accumulation and the turnover of carbohydrates in short photoperiods (8 h light/16 h dark) to compare the nocturnal carbohydrate metabolism in barley and Arabidopsis leaves. Barley primarily accumulated sucrose during the day, followed by small amounts of fructans and starch but did not store fructose and glucose (Figure 1a). At the end of the night, all the sucrose, fructan and starch was consumed (Figure 1a). This confirmed earlier reports stating that soluble storage carbohydrates from the vacuole, respectively sucrose, but not crystalline starch from the chloroplast dominate the carbohydrate supply in the barley leaf (10, 16, 22, 23). As a consequence, barley consumed around five times more carbohydrates from vacuolar sucrose than from starch during the night (Figure 1a). If the conversion of fructans into sucrose before export from the vacuole was considered (7), even seven times more carbohydrates were exported in the form of sucrose from the vacuole than provided from starch degradation in the chloroplast (Figure 1a). In contrast, Arabidopsis accumulated starch in the leaf during the day but no sucrose or glucose and consumed nearly all starch reserves during the night (Figure 1b). As a consequence, Arabidopsis depended almost exclusively on the degradation of chloroplastic starch for carbohydrate supply in the dark (Figure 1b). We then tested if the carbohydrate supply during the night is under control of the circadian clock in barley and Arabidopsis. We measured the depletion of sucrose and the degradation of starch in the leaf of wild-type and *early flowering 3* (*elf3*)-mutant plants of barley (*hvelf3*) (24) and Arabidopsis (*atelf3*). The oscillator in the *elf3*-mutant of both species is severely disturbed as it is arrhythmic in constant light and partly arrested in day/night cycles (24, 25). Depletion of sucrose from the barley leaf did not differ between *hvelf3* and wild-type plants during the night (Figure 1c). However, starch degraded significantly faster in the last half of the night in *hvelf3*-plants compared to wild-type plants (Figure 1d). In Arabidopsis, sucrose levels remained low throughout the night and did not differ between *atelf3* and wild-type plants (Figure 1e). By contrast, the degradation of transitory starch terminated prematurely 2 h before the end of the night in *atelf3* while it continued its linear trend until the end of the night in wild-type plants (Figure 1f). These findings demonstrated that a functional ELF3 was required for the temporal control of starch breakdown at the end of the night in both barley and Arabidopsis, although species-specific differences in the degradation pattern existed. In contrast, the depletion of sucrose from the vacuole in barley was not under the control of ELF3. Sucrose depletion was also not affected by the shortfall of starch degradation in the barley *hvslf3*-mutant at the end of the night (Figure 1c, d). The Arabidopsis *atelf3*-mutant, however, was impaired in starch degradation which represented the primary carbohydrate source during the night. To demonstrate that *atelf3* experienced carbohydrate starvation after the termination of starch degradation before dawn (Figure 1f), we analyzed the expression of the sugar-repressed genes At3g59940 and BRANCHED CHAIN AMINO ACID TRANSFERASE 2 (BCAT2) as starvation reporters (4, 26) (Figure 1g, h). Both transcripts were upregulated 2 h before the end of the 24 h cycle in Arabidopsis *atelf3*-mutant plants but not in wild-type plants (Figure 1g, h). Wild-type plants only increased expression of the starvation markers when they were kept in extended darkness beyond 24 h after dawn (Figure 1g, h). As even small but repetitive reductions of carbohydrate supply at night reduce growth in plants (2–4), we tested if the *elf3*-mutants of barley and Arabidopsis were impaired in biomass accumulation. We found that the *elf3*-mutant of Arabidopsis, but not that of barley, showed a significantly reduced fresh and dry weight in comparison to wild-type plants in the growth chamber and in the greenhouse at different stages of vegetative growth (Figure 2a-c, Supplementary Figure 1). Therefore, our data demonstrates that nocturnal carbohydrate supply for metabolism and growth is determined by the circadian clock component ELF3 in Arabidopsis but not in barley. This explains why only *atelf3*, but not *hvelf3*, reduces biomass accumulation during vegetative growth.

**Figure 1:**
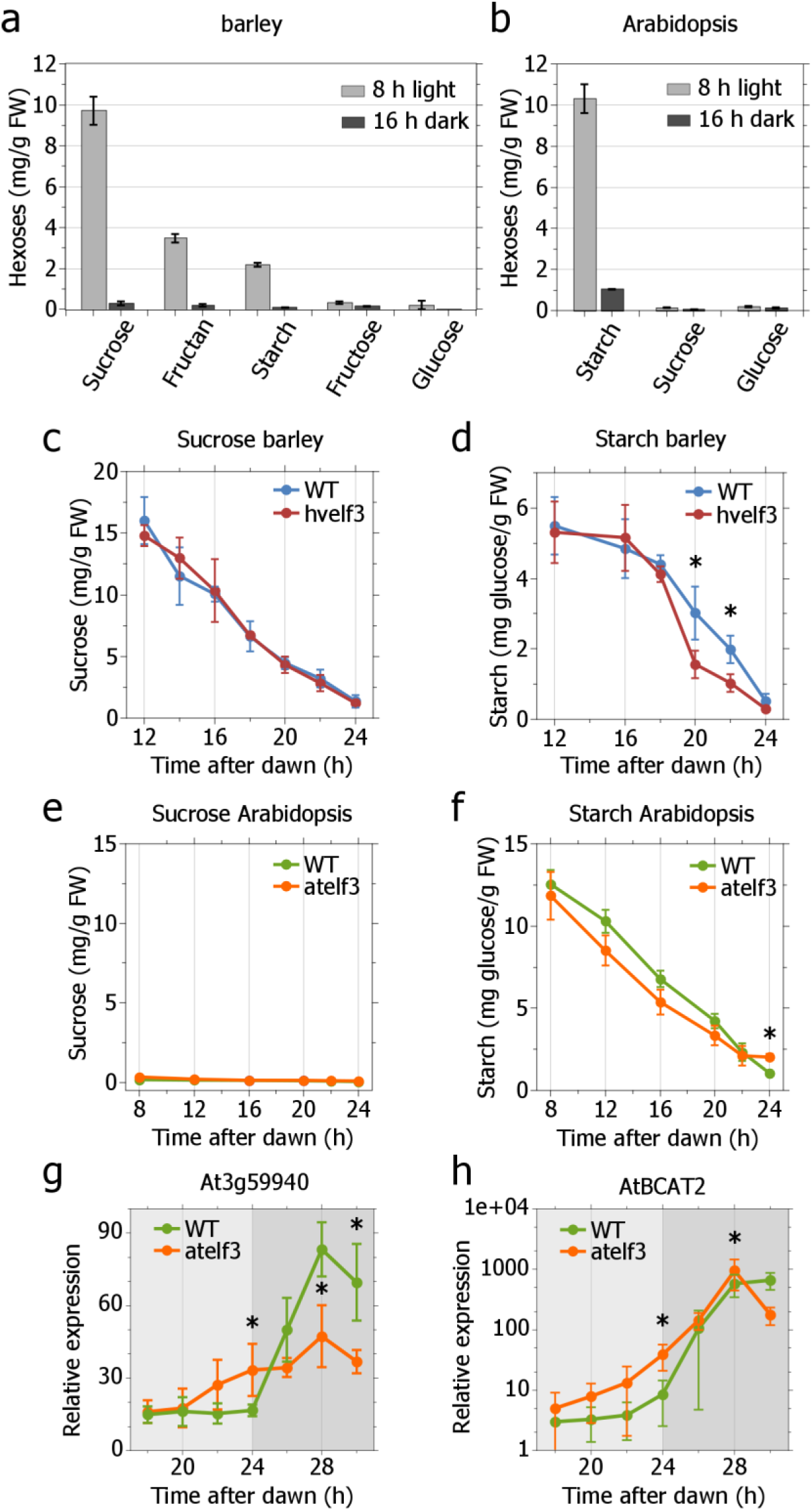
Sucrose depletion dominates carbohydrate supply from the barley leaf during the night and is not controlled by ELF3 while starch degradation dominates carbohydrate supply from the Arabidopsis leaf and requires ELF3 to prevent starvation before dawn. **a**), **b**) Content of carbohydrates in the leaf of a) barley and b) Arabidopsis after the end of the day and the end of the night in 8 h light/16 h dark photoperiods. *N=4 per time point*, **c**), **d**) Content of c) sucrose and d) starch in the leaf of barley *elf3*-mutant and wild-type plants during the night in 12 h light/12 h dark photoperiods. *N=4per time point*, **e**), **f**) Content of e) sucrose and f) starch in the leaf of Arabidopsis *ef3*-mutant and wild-type plants during the night in 8 h light/16 h dark photoperiods. *N=4per time point*, **g**), **h**) Expression of the starvation reporters g) At3g59940 and h) AtBCAT2 in Arabidopsis *elf3*-mutant and wild-type plants before dawn (light grey) and in extended darkness (dark grey). *N=4per time point*. *Significant differences per time point (t-test, p≥0.01) are marked with ^∗^. Error bars are the standard deviation of the mean throughout*.

**Figure 2:**
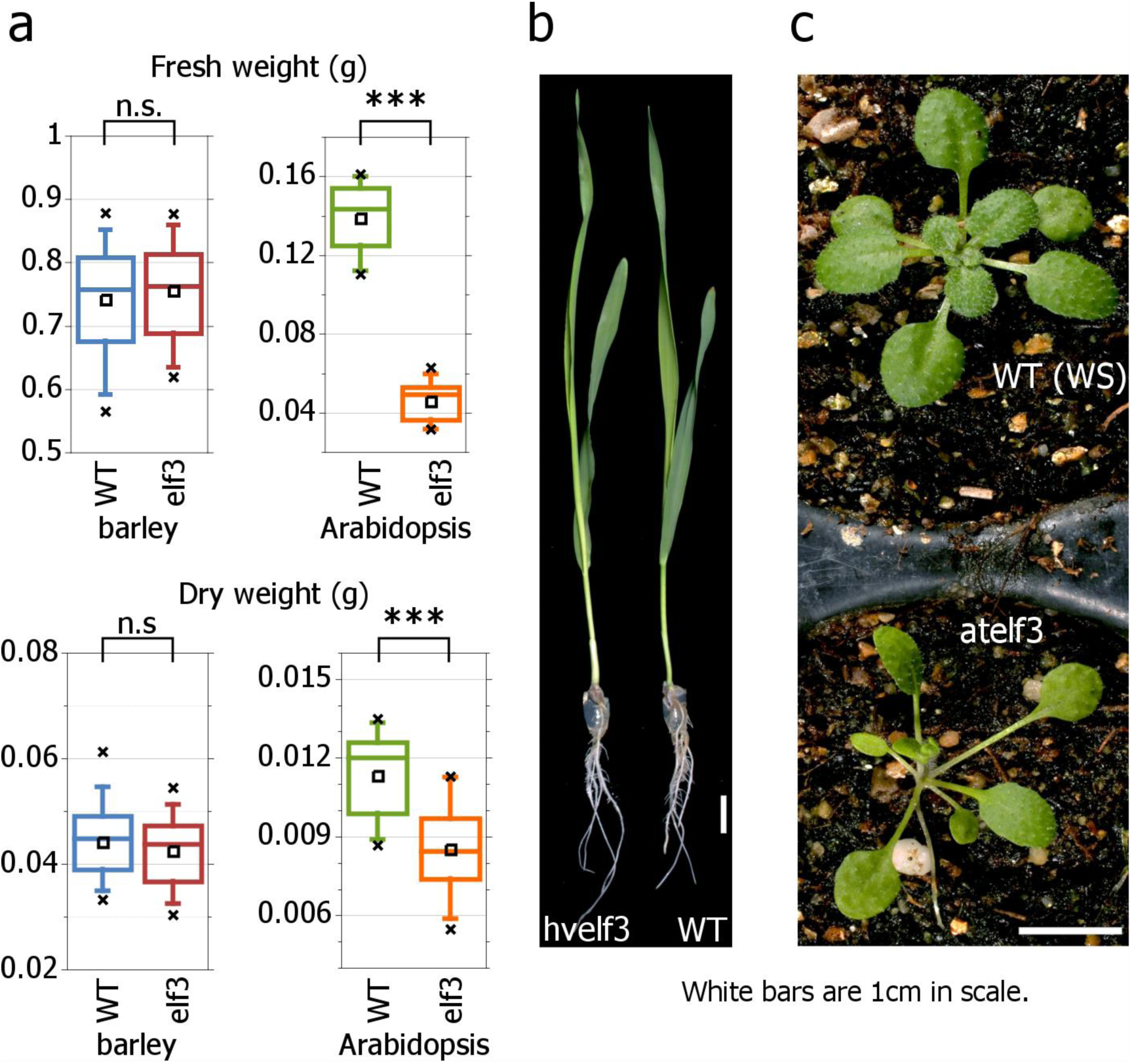
The clock component ELF3 controls biomass accumulation during vegetative growth in Arabidopsis but not in barley. **a**) Fresh and dry weight of *elf3*-mutant and wild-type plants from barley and Arabidopsis during vegetative growth before transition to reproductive growth. *N≥20*. **b**), **c**) Phenotypes of b) barley and c) Arabidopsis *ef3*-mutant and wild-type plants. *Boxes show the 25 and 75% percentile, whiskers denote the 5-95% percentile, the line denotes the median and the square denotes the mean. Significant differences were tested by t-test (p≤0.01). White scale bars represent 1cm*.

### Sucrose depletion from the leaf is controlled by the sucrose content and the night temperature through an exponential kinetics

Sucrose levels in the barley *elf3*-mutant were almost exhausted at the end of the night 24 h after dawn although sucrose depletion was not controlled by ELF3 (Figure 1c). We, therefore, investigated if nocturnal sucrose depletion from the leaf is controlled by a clock-independent mechanism to adjust carbohydrate supply to the length of the night. We grew wild-type plants under cycles of 8 h light/16 h dark and measured the sucrose content over the whole diel cycle. During the night, sucrose depletion was exponential and sucrose levels 24 h after dawn were identical to those 24 h before (Figure 3a). This cycle demonstrated that sucrose depletion is under temporal control. To analyze the adaptability of sucrose depletion to the length of the night, we exposed plants to an unexpected extension of the light period from 8 h to 12 h. This extension increased the sucrose content in the leaf at the end of the light period and enhanced the sucrose depletion rate at night so that levels were comparable between entrained and extended photoperiods at the end of the night 24 h after dawn (Figure 3a). This demonstrated that the sucrose content at the end of the light period relates to the rate of sucrose depletion during the night so that sucrose is almost completely exhausted in the leaf 24 h after dawn, even after unexpected variation of the light period.

**Figure 3:**
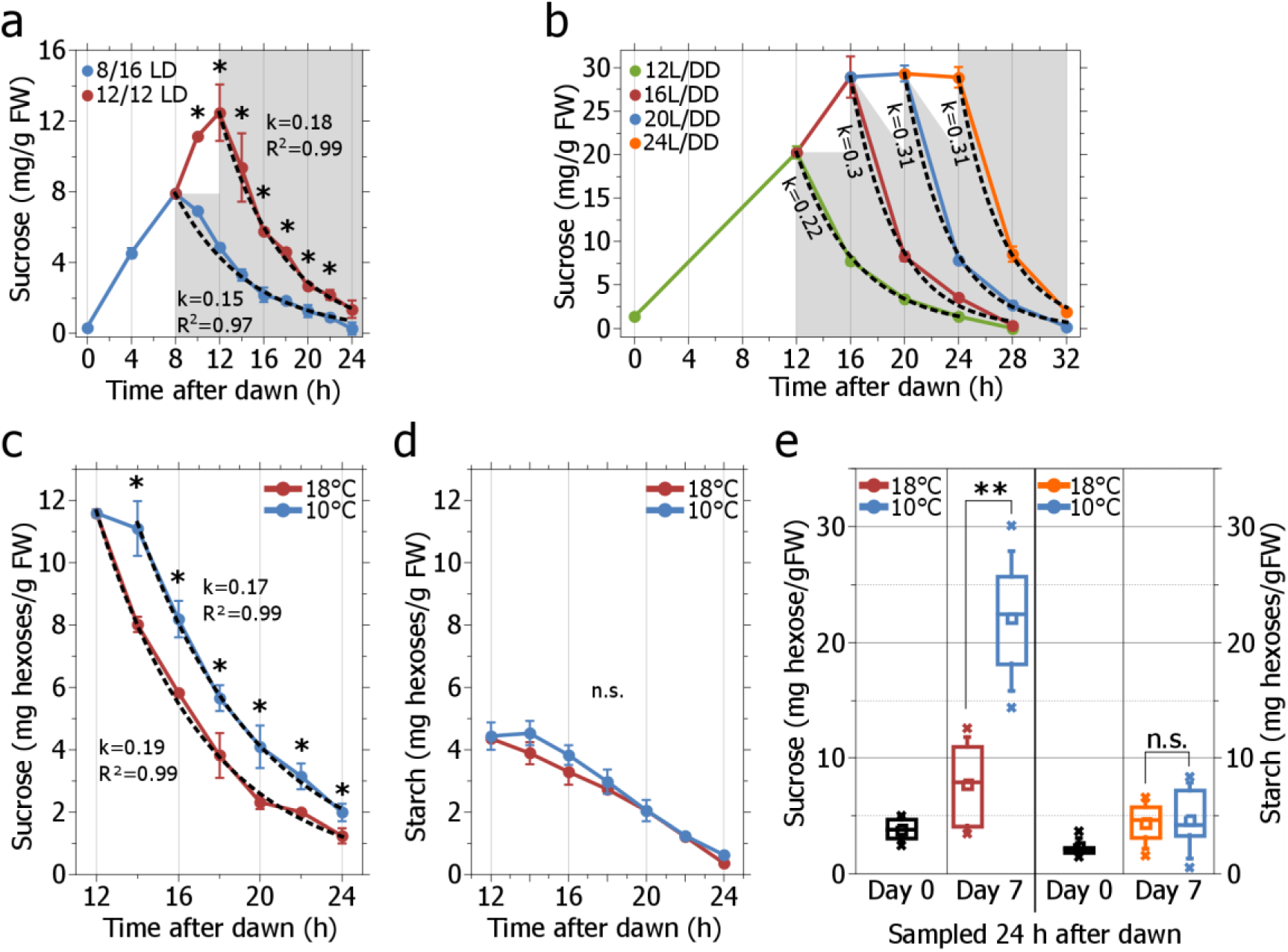
The sucrose content and the temperature control the depletion of sucrose from the barley leaf at night while starch degradation is compensated against low temperature. **a**) Leaf sucrose content in wild-type barley plants entrained to a photoperiod of 8 h light/16 h dark and then kept in 8 h light/16 h dark (8/16 LD, blue) or unexpectedly shifted to 12 h light/12 h dark (12/12 LD, red). *N=4 per time point*, **b**) Leaf sucrose content in wild-type barley plants entrained to 12 h light/12 h dark and then kept in 12 h light (12 L, green) and released to constant darkness (DD) or unexpectedly shifted to an extended light period of 16 h (16 L, red), 20 h (20 L, blue) or 24 h (24 L, orange) before being released to constant darkness (DD). *N=3per time point*, **c**), **d**) Content of c) sucrose and d) starch in the leaf of barley wild-type plants entrained to 12 h light at 20°C and 12 h dark at 18°C before plants were exposed to either an unexpected cool night at 10°C (blue) or kept in the entrained night temperature of 18°C (red). *N=4 per time point*, **e**) Sucrose and starch content at dawn in three week old barley plants grown in 12 h light at 20°C and 12 h dark at 18°C (Day 0) and after seven days of growth at 20°C during the day and either 18°C or 10°C during the night (Day 7). *Significant differences per time point (t-test, p≤0.01) are marked with ^∗^. Error bars are the standard deviation of the mean throughout. Boxes show the 25 and 75% percentile, whiskers denote the 5-95°% percentile, the line denotes the median and the square denotes the mean.*

We modeled the sucrose export from the leaf during the night as a function of the sucrose concentration and fitted the equation of first-order chemical kinetics [suc]_t_ = [suc]_0_ e^-kt^ to the measured data. Chemical kinetics of the first order describes enzyme catalyzed reactions that depend, at constant temperature, only on the substrate concentration. We were able to approximate the sucrose content in the leaf over time based on the sucrose content at the beginning of the night and extracted the depletion constant *k* to quantify the rate of sucrose depletion. When the light period was unexpectedly extended from 8 h to 12 h, the rate of sucrose depletion was 0.03 units higher than under entrained conditions (*k*≈0.18 vs. *k*≈0.15), reflecting the increased sucrose depletion rate that was necessary to deplete a higher amount of sucrose in a shorter night (Figure 3a). To further investigate the adaptability of the sucrose depletion kinetics, we analyzed barley plants shifted from 12 h of light to an unexpected 16, 20, and 24 h period of light and released them to constant darkness (Figure 3b). An unexpected extension of the light period from 12 h to 16 h increased the sucrose content in the leaf and increased the sucrose export rate in the dark for 0.08 units from *k*≈0.22 to *k*≈0.3 (Figure 3b). However, an unexpected extension of the light period from 12 h to 20 h or even 24 h saturated the sucrose content in the leaf and the depletion rate in the dark did not increase much further in comparison to 16 h of light (*k*≈0.3 vs. *k*≈0.31) (Figure 3b). Together, this demonstrated a close relationship between the sucrose content in the leaf at dusk and the depletion rate during the following night.

As the activity of metabolic enzymes is temperature-dependent (27), we tested if the night temperature influences the kinetics of sucrose depletion from the leaf. After an unexpected reduction of the night temperature from entrained 18°C to 10°C at the onset of darkness, the sucrose content in the leaf was significantly higher throughout the night at 10°C and the rate of sucrose depletion during the night was 0.02 units lower at 10°C compared to 18°C (*k*≈0.17 and *k*≈0.19, respectively) (Figure 3c). Therefore, less sucrose was turned over in the leaf during the night at 10°C compared to 18°C. The initial delay in sucrose depletion in the night at 10°C in Figure 3c was probably due to a temperature shock that plants experienced when they were transferred from 20°C to the dark pre-cooled growth chamber of 10°C at the end of the light period. However, even after this initial delay, sucrose depletion rates were lower under 10°C compared to 18°C. This reduction in the sucrose depletion rate in cool nights was comparable to the change observed after an unexpected extension of the day by 4h (change in *k* after temperature drop in Figure 3c: Δ*k*≈0.2, change in *k* after day extension in Figure 3a: Δ*k*≈0.3). In contrast, the starch content was, after a similar initial delay, not significantly different throughout the night between 10°C and 18°C. Consequently, the rate of starch degradation was identical between the cool and the entrained night conditions (Figure 3d). This demonstrated that an internal mechanism was capable to compensate the rate of starch degradation, but not that of sucrose depletion, against a sudden reduction in night temperature. We then investigated how the sucrose depletion and the starch degradation adapted to consecutive cool nights. We grew 3 week old barley wild-type plants for one further week at 20°C during the day and either the entrained 18°C or the cool 10°C during the night. The light intensity during the day was identical for both treatments. At the seventh day, plants grown in the 10°C cool night contained around three times more sucrose in the leaf at the end of the night than those from 18°C (Figure 3e). In contrast, the starch content was not significantly different between both temperature treatments at the end of the night and similar to the onset of the experiment at day 0 (Figure 3e). This demonstrated that the nocturnal sucrose depletion in barley does not adapt to consecutive cool nights and is under environmental and not endogenous control. It also indicated that the effects of consecutive cool nights amplify the incomplete turnover of sucrose in barley as observed for a single cool night in Figure 3c. When taken together, our findings demonstrate that the sucrose content and the night temperature control the depletion of sucrose from the barley leaf in the dark following first-order chemical kinetics. Variation in the sucrose content at the onset of the night alters the depletion rate so that sucrose levels exhaust at dawn, even under unexpected variation of the photoperiod. On the contrary, low night temperatures reduce the sucrose depletion rate, resulting in an incomplete turnover of sucrose during the night. This effect is magnified by consecutive cool nights. Starch degradation, on the other hand, is controlled by an endogenous mechanism in barley that involves the circadian clock component ELF3 and is capable of compensating the degradation rate against low night temperatures.

### Cool night temperatures reduce growth (biomass accumulation) in species predominantly consuming sucrose instead of starch during the night

Pyl *et al*. (2012) (6) identified unused capacity in biosynthetic pathways and processes for growth in Arabidopsis plants in cool nights. They concluded that the carbohydrate supply but not the biochemical pathways involved in growth limits biomass accumulation during nights at low temperature (6). They demonstrated in Arabidopsis that the systemic control of starch degradation through the circadian clock adjusts enzyme activity for starch breakdown to low temperatures so that the rate of starch degradation and, as a consequence, the carbohydrate supply and, subsequently, growth was compensated against cool nights (6). As barley consumes both sucrose and starch during the night but only the degradation of starch was compensated against low night temperature (Figure 3 c-e), we tested if the reduction in sucrose depletion from the barley leaf during cool nights affects growth. We confirmed that growth in Arabidopsis plants is compensated against cool nights of 10°C as fresh weight biomass did not differ between plants grown for one week in cool versus control nights (Figure 4a). However, the fresh weight of barley plants grown in 10°C nights for one week was significantly lower than that of plants grown under control night temperature of 18°C (Figure 4b). Therefore, the reduced sucrose turnover in the barley leaf at low night temperatures correlated with reduced growth. This suggested that low night temperatures reduced carbohydrate supply and, subsequently, nocturnal growth in barley. We then investigated if the sucrose depletion metabolism from barley and its associated growth reduction in cool nights is a domestication-related trait and if it is present in further monocot species. Wild barley, wheat and rice metabolized approximately three times more sucrose than starch during the night (Figure 5a, Supplementary Figure 2). By contrast, *Brachypodium* consumed 1.4-fold more starch than sucrose during the night (sucrose-to-starch ratio: 0.7, Figure 5a, Supplementary Figure 2). Wild barley, rice, and wheat, but not *Brachypodium*, significantly reduced fresh weight after one week of growth in cool nights (Figure 5b). These findings showed that the sucrose depletion metabolism and its association with reduced growth in cool nights is prevalent in several agriculturally important cereal crops. This includes rice, barley and wheat but not all monocot species as nocturnal growth in *Brachypodium* primarily depends on starch instead of sucrose and was compensated against low night temperature. Moreover, the sucrose depletion metabolism is apparently not a domestication trait because it is present in both wild and cultivated barley (Figure 5a, b). When taken together, we observed that the primary carbohydrate store correlated with differences in the growth response of monocot species to cool nights. It is likely that the reduced carbohydrate supply from sucrose depletion in cool nights is responsible for the growth reduction, but the causal link between the major carbohydrate store at night and growth responses to external versus internal cues awaits further investigations.

**Figure 4:**
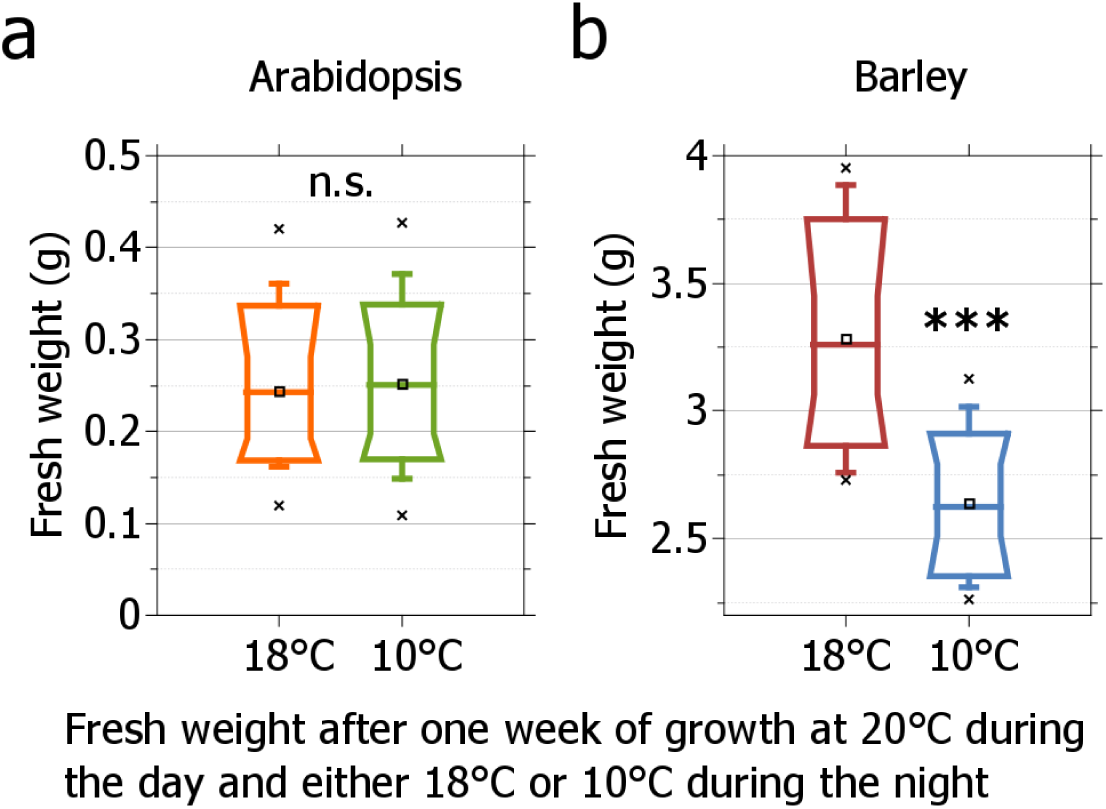
Barley, but not Arabidopsis, reduces biomass accumulation during one week of growth in cool nights. **a**), **b**) Fresh weight of a) Arabidopsis and b) barley wild-type plants after one week of growth in the entrained night temperature of 18°C or cool nights of 10°C. Plants were grown at 20°C during the day in 12h light/ 12 h dark photocycles and the light intensity was identical in both treatments. *Significant differences per time point (t-test, p≤0.0001) are marked with ^∗∗∗^. Boxes show the 10, 25, 75and 90% percentile, whiskers denote the 5-95% percentile, the line denotes the median and the square denotes the mean.*

**Figure 5:**
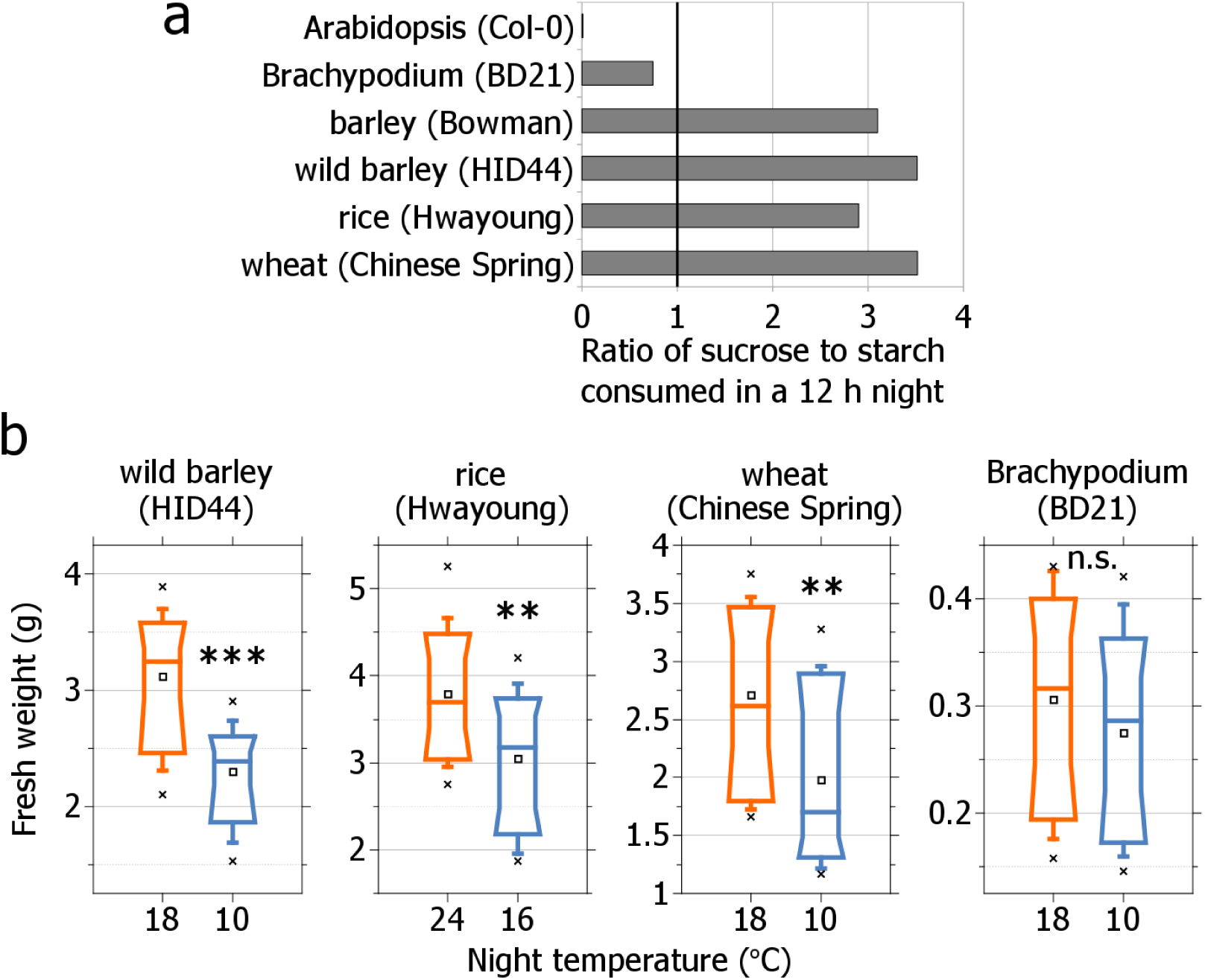
Biomass is reduced during growth in cool nights in several monocot species when they predominantly consume sucrose instead of starch during the night. **a**) Ratio of sucrose to starch consumed during the night in 12 h light/12 h dark photoperiods in several monocot species. A ratio of 1 indicates equal consumption of sucrose and starch during the night. **b**) Fresh weight of species depicted in a) after one week of growth at a low night time temperature compared to the entrained favorable night time temperature for the particular specie. *N=50*. *Significant differences per time point (t-test, p≤0.01) are marked with ^∗^. Boxes show the 10, 25, 75and 90% percentile, whiskers denote the 5-95% percentile, the line denotes the median and the square denotes the mean.*

### Sucrose export from the vacuole catalyzed by SUCROSE TRANSPORTER 2 regulates the sucrose depletion metabolism and affects growth and yield

Finally, we investigated the molecular basis of the sucrose depletion metabolism in cereal crops. Previous studies in barley, wheat and rice demonstrated that the SUCROSE TRANSPORTER 2 (SUT2) is located in the vacuolar membrane and catalyzes the export of sucrose through a proton coupled co-transport (13, 14, 28, 29). Isolation of the *sut2*-mutant in rice provided evidence *in planta* that SUT2 is necessary for the sucrose export from the vacuole into the cytoplasm as *sut2*-mutant plants over-accumulated sucrose in the leaf vacuole at dawn (14). As rice, like barley, depends on the sucrose depletion metabolism for carbohydrate supply at night (Figure 5a, b) and the barley *sut2*-mutant was not available to us, we investigated in rice if SUT2 is involved in the regulation of the nocturnal sucrose depletion metabolism. The sucrose depletion metabolism is based on a positive relationship between the sucrose content in the leaf at dusk and the amount of sucrose depleted in an exponential fashion during the following night (Figure 3a, b). While sucrose depletion was exponential with a constant of *k*≈0.12 in wild-type plants, sucrose depletion in the *sut2*-mutant was rather linear than exponential due to the low depletion constant of *k*≈0.03 (Figure 6a). In addition, despite the nearly two-fold higher sucrose content at dusk, the *sut2*-mutant depleted only around half of the amount of sucrose per Gramm fresh weight (mg/gFW) until dawn than wild-type plants (13mg and 22mg sucrose/gFW, respectively) (Figure 6a). This demonstrated that SUT2 function is required for the exponential depletion of sucrose from the leaf based on the sucrose content. This disturbance of the sucrose depletion metabolism correlated with a strong reduction of growth, biomass and 1000 kernel weight in the *sut2*-mutant (Figure 6b-d). This confirmed that the sucrose export from the leaf vacuole during the night affects growth and yield. The functional redundancy of the SUT2 transporter in rice, barley and wheat (13, 14, 28, 29) and the predominant consumption of sucrose instead of starch in these species during the night (Figure 5a) together with the growth reduction in cool nights (Figure 5b) suggests that the vacuolar sucrose exporter SUT2 is also a key regulator of the sucrose depletion metabolism and growth in barley and wheat. By contrast, Arabidopsis plants, which accumulate almost exclusively starch in the chloroplast during the day but almost no sucrose in the vacuole (Figure 1b), do not express the homologue of OsSUT2, AtSUT4/SUC4, in the leaf (30). This highlights the different regulatory principles that exist in plants for the depletion of sucrose and the degradation of starch in the leaf during the night and their specific effects on growth.

**Figure 6:**
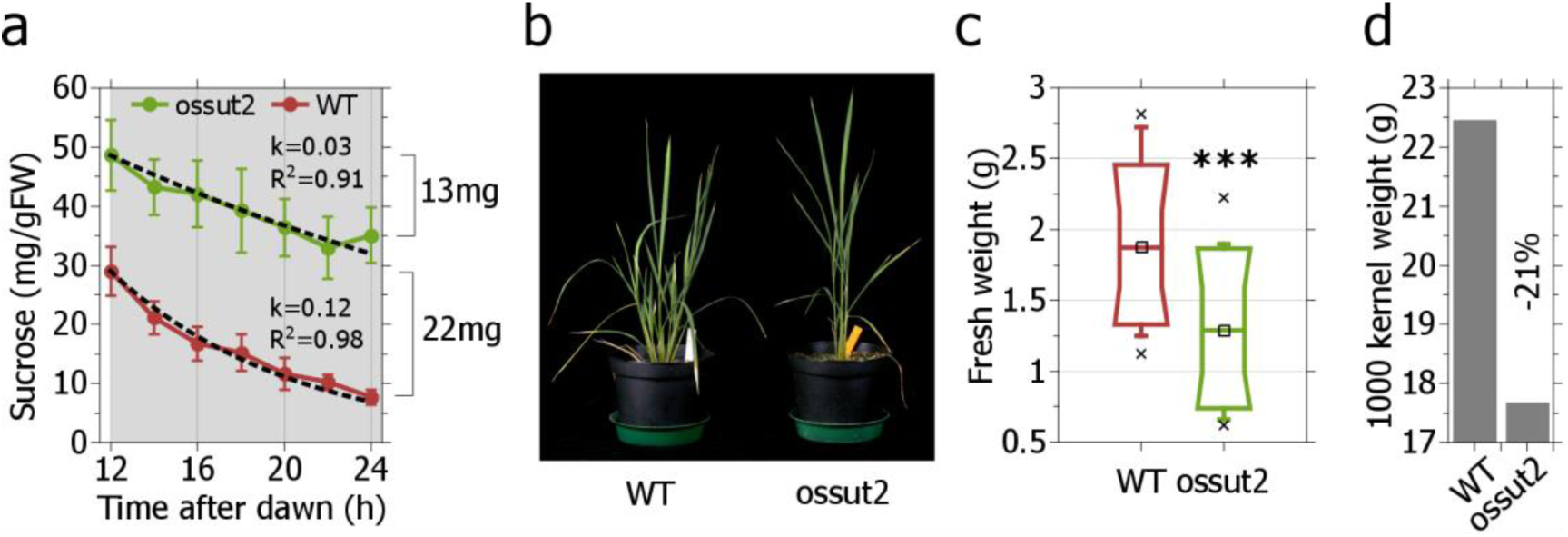
Sucrose export from the vacuole catalyzed by SUCROSE TRANSPORTER 2 (SUT2) in rice plays a key role in the regulation of the sucrose depletion metabolism. **a**) Sucrose depletion from the leaf during a 12 h night of rice *sut2*-mutant and wild-type plants. **b**), **c**), **d**) Loss of SUT2-function reduces b) growth, c) biomass and d) 1000 kernel weight in rice. *Significant differences per time point (t-test, p≤0.01) are marked with ^∗^. Boxes show the 10, 25, 75and 90% percentile, whiskers denote the 5-95% percentile, the line denotes the median and the square denotes the mean.*

### Conclusion

Our results suggest that metabolism and growth in plants at night is determined by the predominant nocturnal carbohydrate stores of starch and sucrose. The nocturnal starch turnover is controlled by the circadian clock, whereas sucrose depletion is determined by vacuolar sucrose concentrations at the end of the day and night-time temperatures. This endogenous versus external regulation of carbohydrate supply correlates with variation in biomass accumulation between sucrose and starch storing species. In this context, it remains to be understood why different species either use starch or sucrose or a combination of both as transient storage carbohydrates for the night. The starch storing mechanism might buffer growth against environmental fluctuations, but the sucrose depletion metabolism may offer a growth advantage over the starch degradation metabolism because the carbohydrate supply from low molecular sucrose is energetically advantageous compared to the synthesis and the degradation of high molecular starch. However, the nocturnal sucrose export and the subsequent growth are susceptible to cool nights. A “hybrid” system with sucrose and starch as seen in *Brachypodium* may combine the benefits from sucrose-driven growth under favorable and starch-driven growth under unfavorable conditions. Barley, wheat and rice might accumulate little amounts of starch during the day to mitigate a growth reduction in cool nights.

We demonstrate that the physiological control of growth in important cereal crops cannot be resolved in the current model species Arabidopsis and *Brachypodium*, despite the conservation of clock components in crop plants (17, 20, 21, 31). We propose that the sucrose depletion metabolism gives new insight into the physiology of growth and yield in cereal crops. In addition, the monocot lineage with variation in carbohydrate storage provides the opportunity to study the genetics, evolution and adaptive significance of carbohydrate use and growth in plants. A mechanistic understanding of growth as controlled by sucrose versus starch is important for the adaptation of crops to climate change.

## Acknowledgments

This work was supported by the Max Planck Society through an International Max Planck Research School fellowship to L.M.M., by an ERASysApp project (“CropClock”), a trilateral DFG grant (“Circadian clock and stress adaptation in barley”) and by the Excellence Cluster EXC1028.

## Author contributions

L.M.M. conceived the study and conducted the experiments; L.M.M. and M.v.K. designed the experiments; L.G. contributed experimental data; L.M.M., M.v.K., A.P.M.W., and S.J.D. analyzed the data; J-S.J. contributed new material; L.M.M. and M.v.K. prepared the manuscript.

The authors declare no competing financial interests.

Correspondence and requests for materials should be addressed to L.M.M. or M.v.K.

## Materials and Methods

### Plant material and growth conditions

The spring cultivar Bowman (BW) was used as the wild-type barley. The introgression line BW290, which carries an introgression of the *eam8.k* allele in the background of Bowman, was denoted by *hvelf3*. The *eam8.k* allele is characterized by a base-pair mutation leading to a premature stop codon in *HvELF3*, which is orthologous to *ELF3* in Arabidopsis (24). For Arabidopsis, the *elf3*-*4* mutant in the Wassilewskija (WS) background was used (32). Both barley and Arabidopsis were grown in growth chambers at 450 μmol m^−2^ s^−1^ photon flux density (PPFD) at 20°C during the day and 18°C during the night as standard growth conditions. For the cool night treatment, standard growth conditions were applied during the day and only the nighttime temperature was altered to 10°C. The significant difference in biomass between *hvelf3* and wild-type plants was confirmed in a temperature-controlled greenhouse (20°C during the day, 18°C during the night). Both barley and Arabidopsis seeds were directly sown into soil (Einheitserde) and stratified for 4 days at 4°C in the dark.

The rice mutant *ossut2* carried a T-DNA insertion in the fifth exon of the *SUT2* gene in the Hwayong background (14). The rice seeds were directly sown into potting soil (Einheitserde) and cultured in growth chambers under 12 h light at 28°C and 350–450 μmol m^−2^ s^−1^ photon flux density (PPFD) and 12 h dark at 25°C.

### Diel/circadian sampling

Barley and rice plants were harvested during vegetative growth and grown to the three-leaf stage. The whole second leaf of two plants was cut at the base and placed into a 2 ml Eppendorf tube (200-300 mg) as one biological replicate. For Arabidopsis, plants were grown for 4 weeks before complete rosettes from at least 4 different plants were pooled to one biological replicate (100-200 mg). At least 4 different biological replicates were sampled per time point, genotype and plant species and snap frozen in liquid nitrogen. The time points of sampling were applied as given in the respective diagrams of the individual experiments.

### Biomass measurements

Biomass was measured as the fresh weight of the shoot on an accuracy balance directly after cutting from the rootstock. Individual shoots were collected, oven dried and measured for dry weight. At least 20 biological replicates were measured per genotype for the biomass analysis. Arabidopsis plants were harvested before the first bud became visible, while the development of the monocot species was established by dissecting the shoot apical meristem and scoring the development of the main shoot apex as described in (33). This ensured that biomass measurements captured the vegetative and not the reproductive growth.

### Measurement of sugars and starch

Frozen samples were ground to powder using milling beads. The soluble sugar fraction was separated from the insoluble starch fraction by boiling the samples twice in 1 ml of 80% (vol/vol) ethanol at 85°C for 30 min. Chlorophyll was precipitated from the soluble fraction by adding 0.5 ml of chloroform and 1 ml of water. The clear alcoholic extract was concentrated in a vacuum concentrator and resuspended in sterile water. The content of sucrose, glucose, fructose, and maltose was measured using a photometer at 340 nm wavelength as NADPH absorbance in a glucose-6-phosphate dehydrogenase (G6PDH)-dependent assay and quantified against a glucose standard curve using glucose/sucrose enzyme kits from r-biopharm (Cat. No. 11 113 950 035).

To measure fructan content, the resuspended concentrate was hydrolyzed with 37% HCl for 20 min at 80°C, then neutralized with 7.67 M NaOH and measured as fructose against the fructose background using an enzyme kit to measure fructose (r-biopharm, Cat. No. 10 139 106 35).

For the measurement of starch, the insoluble fraction was boiled in KOH at 95°C for 45 min, neutralized with acetic acid (HAc), digested to glucose by *α* -amylase and amyloglucosidase at room temperature overnight, and quantified as glucose.

### Equation of first-order enzyme kinetics

The equation [suc]_t_ = [suc]_0_ e^-kt^ was fitted to the sucrose content at night to describe the depletion kinetics. [suc]_t_ is the sucrose concentration in the leaf at any given time *t* during the night, [suc]_0_ is the initial sucrose concentration at the beginning of the night, and *k* is the depletion constant, which is a measure of the depletion rate.

Differences between treatments or genotypes were analyzed via a t-test (p≤0.01). The number of biological replicates was at least 4 for one time point in the time series data and at least 20 for biomass measurements.

### Transcript Analysis

Five whole rosettes were sampled and pooled per biological replicate, for a total of 4 biological replicates per time point. Extraction of RNA, reverse transcription and qRT-PCR were then performed as previously described (Digel et al., 2015). Briefly, RNA was extracted using the TRIZOL reagent (Thermo Fisher Scientific) according to the manufacturer’s instructions. The DNase treated RNA was transcribed into first strand cDNA using Superscript II reverse transcriptase (Thermo Fisher Scientific) in accordance to the manufacturer’s instructions. Two technical replicates were performed for each sample and absolute quantification was based on a titration curve of cloned fragments for each target gene. All qRT-PCR reactions were performed on a LightCycler 480 (Roche; Software version 1.5) using the following primers: AtBCAT2 FW 5’-GTACAACGCACAAG CTGCAT-3’, RV 5’-ACCCGAGATTATCCCAATCC-3’; At3g59940 FW 5’-TGGAACGATGATGGTGAAGA-3’, RV 5’-AACCAGAGGGAGTGTTGACG-3’. The geometric mean of AtPP2A (FW 5’-CAGCAACGAATTGTGTTTGG-3’, RV 5’-AAATACGCCCAACGAACAAA-3’) and AtPEX4 (FW 5’-TTACGAAGGCGGTGTTTTTC-3’, RV 5’-GGCGAGGCGTGTATACATTT-3’) absolute expression was used to calculate relative gene expression values.

## Supplementary Figures

**Supplementary Figure 1:**
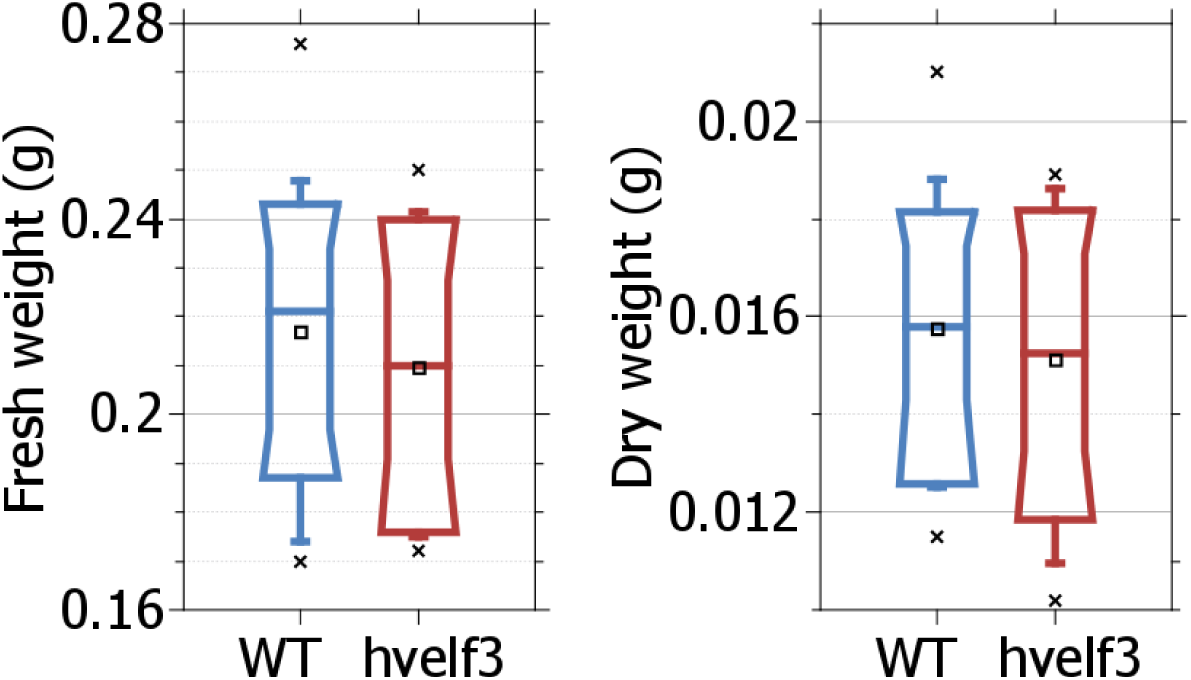
Fresh weight and dry weight of barley *elf3*-mutant and wild-type plants grown in the greenhouse in early summer one week after emergence from the soil. *N=30*.

**Supplementary Figure 2:**
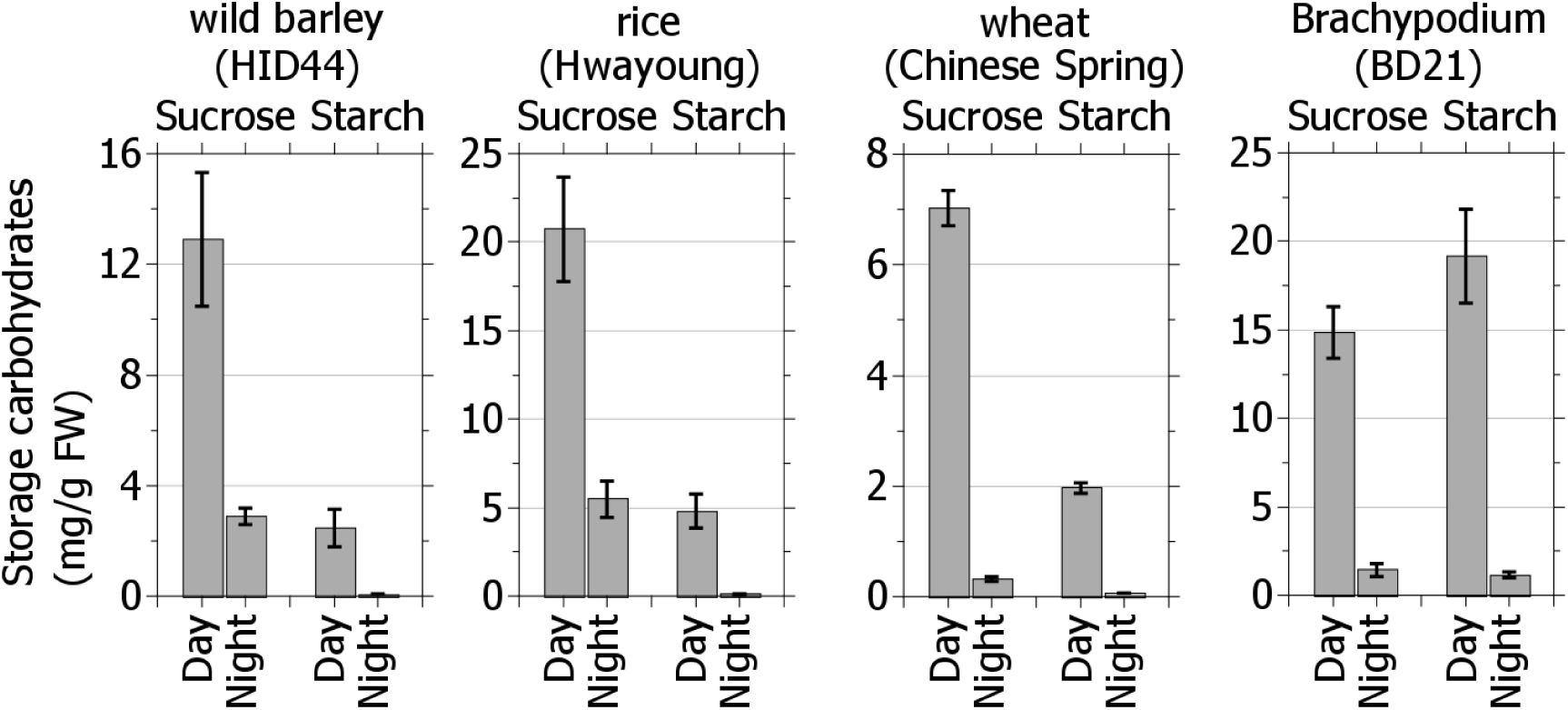
Sucrose and starch content at the end of a 12 h day and a 12 h night in several monocot species. *N=5per time point*.

